# Airway succinate chemosensing induces CFTR-dependent anion secretion and mucus clearance which is impaired in cystic fibrosis

**DOI:** 10.1101/2024.03.26.586799

**Authors:** Tábata Apablaza, Marisol Barros-Poblete, Livia Delpiano, Sandra Villanueva, Anita Guequen, Bárbara Tapia-Balladares, Iram Haq, Felipe Tribiños, Sebastián Hernández-Rivas, Bernard Verdon, Matthew G.S. Biggart, Yenniffer Sánchez, Christopher Ward, B Dnate’ Baxter, Diego Restrepo, Isabel Cornejo, Robert Tarran, Marcelo A. Catalán, Michael A. Gray, Carlos A. Flores

## Abstract

The respiratory tract possesses a highly regulated innate defense system which includes efficient cilia-mediated mucus transport or mucociliary clearance (MCC). This essential process relies on appropriate hydration of airway surfaces which is controlled by a blend of transepithelial sodium and liquid absorption via the epithelial sodium channel (ENaC), and anion and liquid secretion, primarily regulated by the cystic fibrosis transmembrane conductance regulator (CFTR) channel. MCC is tightly regulated by second messenger signalling pathways. Succinate is derived from parasites, microorganisms and inflammatory cells, and its concentration increases in the airway surface liquid (ASL) during infections. Increases in ASL succinate activates the G-protein coupled succinate receptor (SUCNR1), which acts as a succinate sensor. Here, we tested the hypothesis that succinate signalling was linked to CFTR activity, ASL hydration and increased MCC.

We observed that SUCNR1 activation stimulated anion secretion, increased mucus transport and induced bronchoconstriction in mouse airways. In parallel, stimulation of human bronchial epithelial cells (HBEC) with succinate activated anion secretion and increased ASL height. All functions activated by succinate/SUCNR1 were impeded when working with tissues and cells isolated from animal models or individuals affected cystic fibrosis (CF) or when CFTR was inhibited. Moreover, when HBECs derived from ΔF508 individuals were incubated with the triple drug combination of elexacaftor/tezacaftor/ivacaftor (ETI), succinate-induced anion secretion was restored, confirming the tight relationship between SUCNR1 signalling and CFTR function. Our results identify a novel activation pathway for CFTR that participates in the defence response of the airways, which is defective in CF. We propose that succinate acts as a danger molecule that alerts the airways to the presence of pathogens leading to a flushing out of the airways.

## INTRODUCTION

Cystic fibrosis (CF) is a human disease characterized by recurrent lung infections and intermittent exacerbations that ultimately lead to lung function decline. Even though, successful CFTR modulator therapies such as the triple combination of small molecules ETI have restored up to 50% of CFTR function in around 90% of individuals that bear at least one mutant ΔF508 *CFTR* allele ^1,2^, lung infections and inflammation still persist, indicating that the disease is more complex than previously thought ^3,4^.

Substantial evidence indicates that CF presents with defects of the immune system, including hyperinflammatory responses and defective pathogen airway clearance, that negatively affect lung function ^5,6 7^. For example, abnormal increases in inflammatory cytokines such as interleukin-17 (IL-17) and tumor necrosis factor alpha (TNF-α), stimulate the production of mucins that, in the CF airway environment, are inadequately processed, generating poorly hydrated and sticky mucus that occludes the airways and acts as a reservoir for pathogenic bacteria ^8,9 10^. Additionally, animal models with total or cell specific *Cftr* deletion demonstrate impaired immune cell function, as well as abnormal cytokine and immunoglobulin production ^11–14^, indicating that CFTR participates in the immune response.

SUCNR1 is a G-protein-coupled receptor that activates the transient receptor potential channel TRPM5, a non-selective cation channel, whose only known endogenous agonist is succinate, a molecule that can modulate both pro- and anti-inflammatory functions ^15,16^. Succinate and its receptor have been under intense scrutiny in the immunological and metabolic fields, but their role in lung function and airway disease pathogenesis is limited. Detailed explorations of gene expression have determined that brush cells of in murine airways express both *Sucnr1* and *Trpm5* ^17,18^. Brush cells are involved in immuno-surveillance through sensors such as SUCNR1 and the bitter taste receptors TAS2R ^19,20^. These chemosensors can activate MCC in the airways through cholinergic signalling ^21,22^. Even though, brush cells participate in the development of allergic responses and neurogenic inflammation, the involvement of the SUCNR1 has not been studied in airway disease ^23,24^.

Unlike mice that express SUCNR1 in brush cells, humans express the SUCNR1 in a different type of airway cell, the pulmonary neuroendocrine cell (PNEC) ^25^, a rare cell type that participates in the heightened response to allergens in asthmatic patients ^26^. In addition, succinate is increased in the bronchoalveolar lavage of CF patients, favouring *Pseudomonas aeruginosa* colonization ^27^. Altogether, these data suggests that SUCNR1 signalling might be altered in airway diseases, such as CF, but this has never been studied in humans. In our present study we aimed to investigate whether SUCNR1 signalling activates CFTR and whether SUCNR1-dependent functions are impaired in CF cellular and animal models, and whether this signalling is restored after ETI treatment.

## RESULTS

### TRPM5+ brush cells are expressed in the proximal trachea of the mouse and induce anion secretion through the SUCNR1 signalling cascade

Using commercially available anti-SUCNR1 and anti-TRPM5 antibodies, we were unable to detect SUCNR1 or TRPM5 positive cells in mouse airways. Nevertheless, *Sucnr1* transcripts were exclusively expressed in *Trpm5*+ cells derived from mouse tracheas ^17^. Using lung tissues from the TRPM5-eGFP reporter mice we found that TRPM5+ cells were present only in mouse tracheas (Supplemental Figure 1A). Next, we tested several dicarboxylates for their effects on ion transport in ex-vivo airway tissue. We observed that only succinate induced an increase in anion secretion with an EC_50_ of 0.48 mM (Figure 1A).

**Figure 1.**
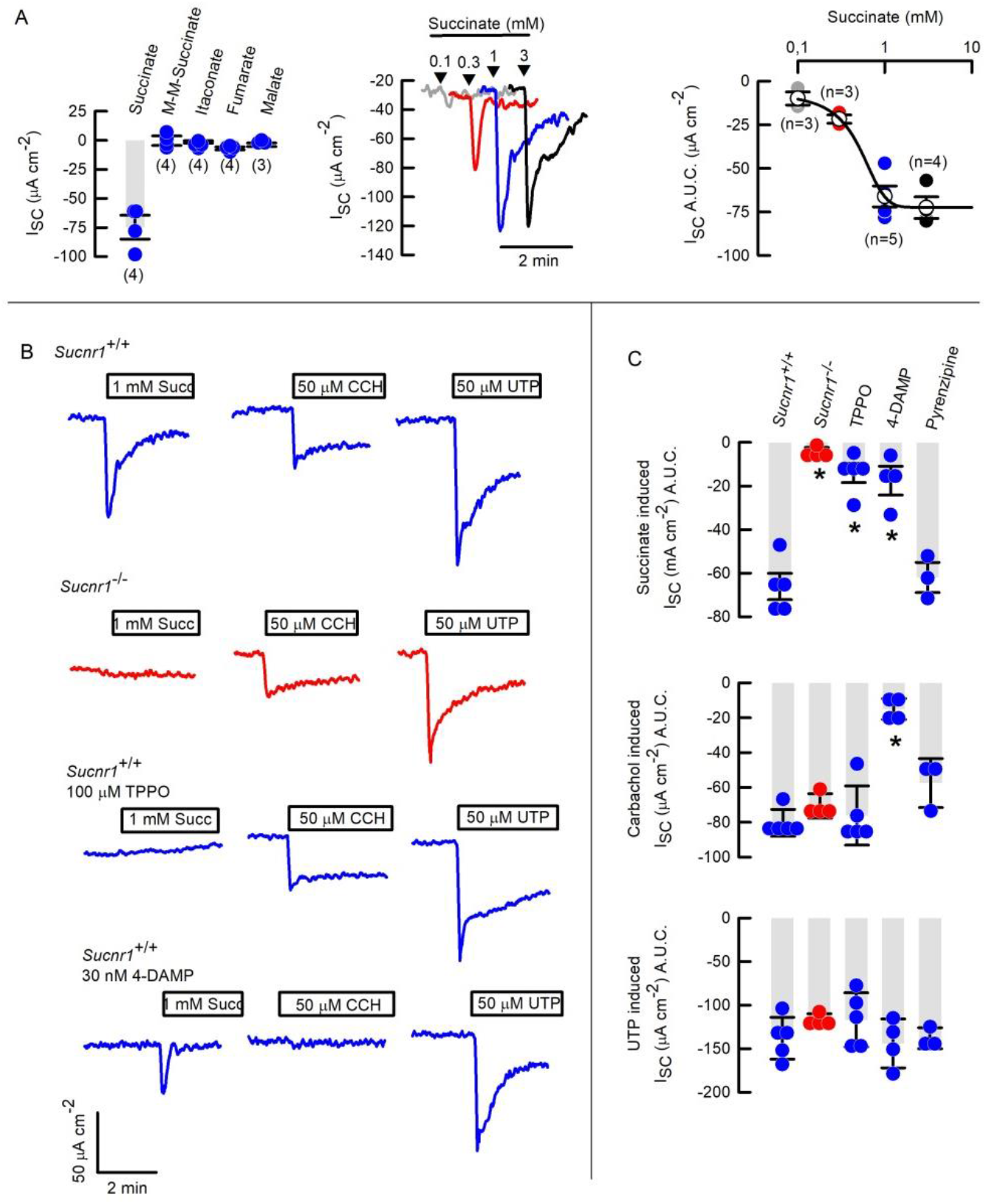
Succinate induces anion secretion through a mechanism that involves SUCNR1, TRPM5 channels and M3 muscarinic receptors in the mouse trachea. (**A**) Summary of the changes in short-circuit current (I_SC_) in mouse trachea in response to the indicated di-carboxylates, representative traces of I_SC_ in mouse tracheas stimulated with increasing succinate concentrations and a summary of succinate-induced I_SC_ in mouse tracheas. (**B**) Representative trace of succinate-, CCh- and UTP-induced anion secretion in wild type, *Sucnr1*^-/-^ tracheas and wild type tracheas incubated with TPPO and 4-DAMP. (**C**) Summary of succinate-induced, CCh-induced and UTP-induced anion secretion in mouse tracheas treated with TPPO, 4-DAMP or pirenzipine; values are Mean ± S.E.M.; n=3-5 per group.* indicates p<0.05, ANOVA on Ranks.

To better understand the signalling events involved in the succinate-induced anion secretion we performed a 3-step Ussing chamber protocol where the SUCNR1 receptor was first activated with 1 mM succinate, which was then followed by stimulation of muscarinic receptors with 50 µM carbachol and finally purinergic signalling with 50 µM UTP, all in the same tissue. As observed in Figure 1B-C, the three agonists induced robust secretory responses. To test if SUCNR1 was involved in the secretory response we generated a *Sucnr1*^-/-^ mouse (Supplemental Figure 2). Succinate-induced anion secretion was absent in tracheas of these mice. In contrast, *Sucnr1*^-/-^ mice exhibited unaltered carbachol and UTP responses Figure 1B-C. Additionally, the response to denatonium, an agonist of TAS2R, was unaltered in *Sucnr1*^-/-^ mice (Supplemental Figure 3A). We then used U732122 to inhibit phospholipase beta2 (PLC-β2) or TPPO to inhibit TRPM5, both key steps in the signal cascade of taste receptors ^22^.These compounds both fully prevented succinate-induced anion secretion (Supplemental Figure 3B and Figure 1B-C).

Brush cells are scarce and do not express anion channels such as CFTR or the calcium-activated anion channel TMEM16A ^17^. It is therefore likely that brush cells do not directly mediate anion secretion. Brush cells are known to synthesize, and store acetylcholine, which can be released upon activation of receptors, including SUCNR1, to induce a muscarinic receptor-dependent activation of neighbouring cells ^22^. Thus, we asked if succinate-induced anion secretion was dependent on this pathway. Accordingly, we tested their involvement by incubating mouse tracheas with pyrenzipine (that preferentially inhibits M1 muscarinic receptors) and dimethyl-4-diphenylacetoxypiperidinium iodide (4-DAMP, an M3 receptor antagonist). As observed in Figure 1B-C, 4-DAMP fully blocked the succinate effect, while pyrenzipine had no effect on either succinate- or carbachol-induced secretion (Figure 1C), indicating that the pro-secretory effect of succinate was transduced by M3-coupled signalling.

### SUCNR1 activation increases mucus transport dependent on bicarbonate secretion, and generates bronchoconstriction in mouse trachea

We observed that succinate induced an increase in particle track speed, an indirect measurement of ciliary beating activity, and therefore MCC. We confirmed that the increase in particle track speed was fully prevented by 4-DAMP (Supplemental Figure 3C-D), as previously demonstrated ^21^. Next, we investigated in more detail the effect of succinate on mucus transport using different protocols, as depicted in Supplemental Figure 4. Using protocol 1 (Supplemental Figure 4B), we observed that tracheas incubated in succinate exhibited faster mucus speed than controls (Supplemental Figure 4C). However, due to the high variability in these experiments the protocol was further adapted to study changes in mucus transport in the same tissue sample (Supplemental Figure 4D). As shown in Figure 2A, it was possible to observe an increase in mucus speed after succinate addition to wild type (Supplemental Video 1) but not *Sucnr1*^-/-^ tracheas (Supplemental Video 2). Mock addition of vehicle to wild type tracheas was also ineffective at stimulating mucus clearance (Supplemental Figure 4E). We observed a significant decrease in mucus velocity in the *Sucnr1*^-/-^ tissues. However, this reduction in MCC was not caused by succinate, since we observed the same effects in vehicle-treated *Sucnr1*^-/-^ tracheas (Supplemental Fig 4E). Finally, pre incubation of tissues with 4-DAMP (protocol 3, Supplemental Figure 4F) fully abolished succinate-dependent increases in mucus transport (Figure 2A).

**Figure 2.**
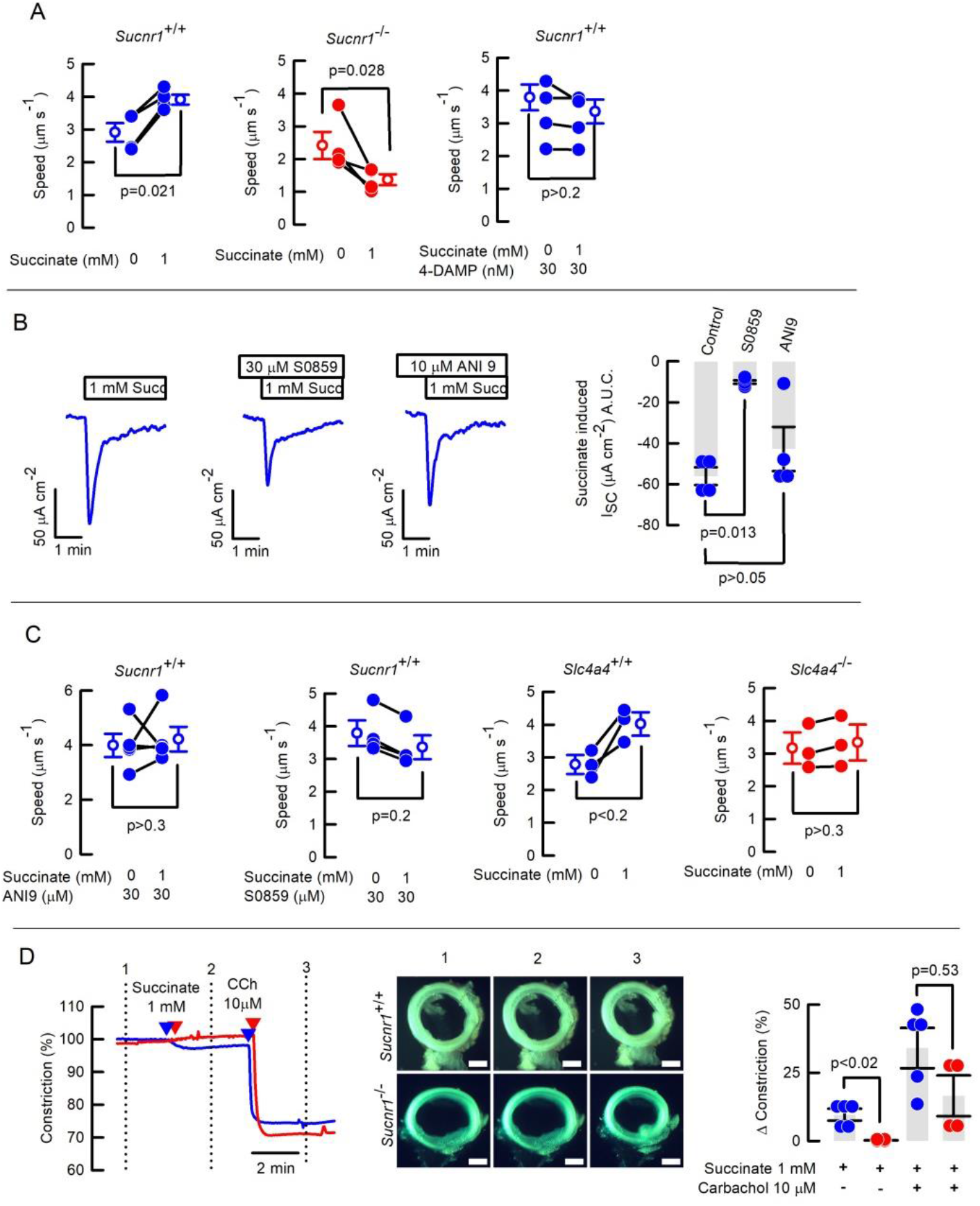
Succinate induces bicarbonate secretion, increased mucus clearance, and induces bronchoconstriction in mouse airways. (**A**) Mucus transport speed before and after succinate in wild type, *Sucnr1*^-/-^ and wild type tracheas treated with 30 nM 4-DAMP. Values are Mean ± S.E.M.; n=4 for each group; Rank-sum test. (**B**) Representative traces for succinate induced I_SC_ in control, after 30 µM ANI9 of 30 µM S0859 and summary of succinate-induced ΔI_SC_, values are Mean ± S.E.M.; n=4 for each group; ANOVA on Ranks. (**C**) Succinate-induced changes in mucus transport speed was determined in wild type mice after 30 µM ANI9 or 30 µM S0859 or in *Slc4a4*^+/+^and *Slc4a4*^-/-^ tracheas. Values are Mean ± S.E.M.; n=4 for each group; Rank-sum test. (**D**) Representative traces of bronchoconstriction in wild type (blue line) or *Sucnr1*^-/-^ (red line) tracheal rings after 1mM succinate and 10 µM carbachol (CCh) addition followed by representative images of tracheal rings; bars = 500 µm. Numbers indicate the moment where corresponding images where taken. Summary of bronchoconstriction experiments, blue dots for wild type and red dots for *Sucnr1*^-/-^. Bars are values are Mean ± S.E.M.; n=5 for each group; Rank-sum test.

Reductions in anion secretion profoundly impair MCC, as observed in CF, where reductions in Cl^-^ and HCO_3_^-^ secretion lead to mucostasis ^28^. On the other hand, an increase in the volume of ASL increases MCC ^29^. We used different inhibitors and animal models to dissect the contribution of Cl- and HCO3-anion transport to the succinate response. Ussing chamber experiments determined that succinate mainly induced HCO_3_^-^ secretion, since the magnitude of the current was significantly reduced by blocking the basolateral Na^+^/HCO_3_^-^ co-transporter SLC4A4 with the inhibitor S0859 (Figure 2B). In contrast, ANI9 which blocks apical TMEM16A channels was ineffective (Figure 2B). We previously determined that particle track speed is significantly reduced when HCO_3_^-^ secretion is decreased ^30^, thus we tested the effect of the inhibitors on mucus transport. Using the pre-incubation protocol (Supplemental Figure 4F), we observed that succinate was unable to increase mucus transport when the tissues were stimulated in the presence of S0859 or ANI9 (Figure 2C). Nevertheless, we observed that ANI9 precipitated in the alcian blue buffer (Supplemental Figure 4G), so we can’t discard the possibility that ANI9 was not effectively altering mucus transport by mechanisms, other than by blocking TMEM16A. To further confirm our observation that the succinate-induced mucus transport increase was dependent on HCO_3_^-^ secretion we used the *Slc4a4*^-/-^ mouse that displays a severe reduction of HCO_3_^-^ secretion but unaltered Cl^-^ secretion ^30^. Using protocol 2 we observed that mucus transport was unaffected by succinate addition in *Slc4a4*^-/-^ tracheas, confirming that bicarbonate secretion was necessary for the increase in mucus transport after SUCNR1 activation (Figure 2C).

Acetylcholine is a known bronchoconstrictor ^31^, and since succinate acts through acetylcholine paracrinally, we tested whether succinate triggered smooth muscle constriction. Using tracheal rings, we observed that succinate induced bronchoconstriction that was not observed in *Sucnr1*^-/-^ tissues (Figure 2D) or when the succinate analogue M-M-succinate was used (Supplemental Figure 5A-B). The effect of succinate was also prevented using either U73122 or TPPO (Supplemental Fig 5C-D), indicating that signalling arising from brush cells triggered bronchoconstriction. Finally, using the data for the changes in bronchoconstriction (Δ constriction), we calculated that succinate caused a 16.9 ± 2.6% reduction in tracheal airflow (Supplemental Fig 5E).

### The M3 receptor is highly expressed in the secretory cells and co-localize with SLC4A4 transporters in the mouse airways

We found that M3 receptors were highly expressed in both surface epithelia and submucosal glands of the proximal airways, unlike the respiratory epithelia where no signal was detected (Supplemental Fig 6A). Functional studies suggested that M3 receptors were expressed in murine airway epithelial ciliated cells ^21,22^. Nevertheless, we found that M3 receptors were highly expressed in club cell 10 positive (CC10+) secretory cells, rather than in acetylated tubulin positive (Ac-tub+) ciliated cells in mouse trachea (Figure 3A). In accordance with our electrophysiological results, the SLC4A4 co-transporter co-localized with M3 receptor, indicating a straight coupling of cholinergic signalling with HCO_3_^-^ secretory machinery in the mouse trachea (Figure 3B). Additionally, we observed that 46% of TRPM5+ cells also showed M3 staining (Figure 3C and Supplemental Figure 6B).

**Figure 3.**
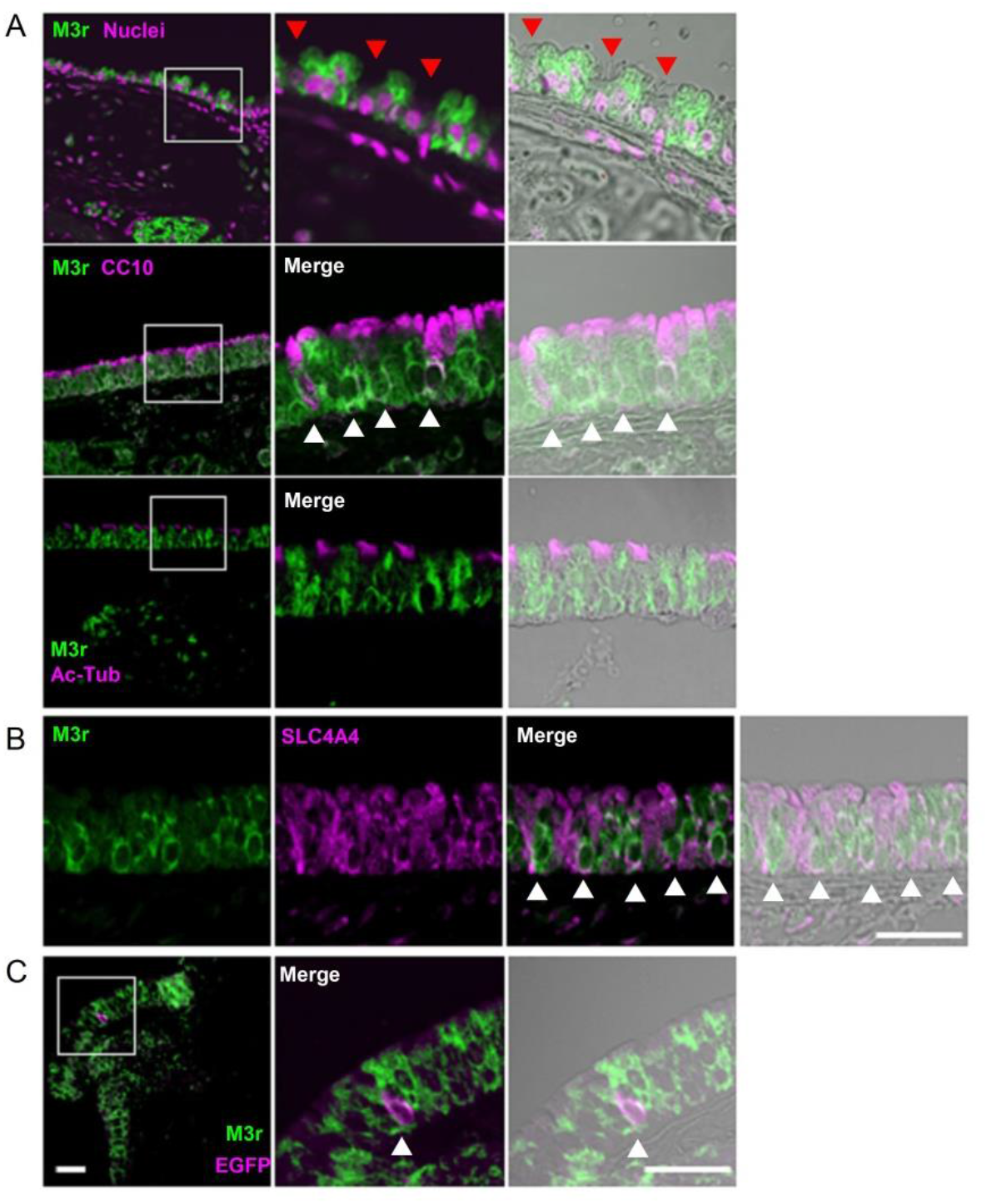
Muscarinic M3 receptors (M3r) are co-expressed with the SLC4A4 co-transporter in mouse airway epithelium. **(A)** M3 muscarinic receptor staining (M3r) is excluded from ciliated cells as indicated by the red arrowheads and acetylated tubulin staining (Ac-Tub), but co-localizes with CC10 staining in secretory or club cells indicated by white arrowheads. **(B)** SLC4A4 co-localizes with M3r staining as indicated with the white arrow heads. **(C)** EGFP positive cell showing M3r staining. Representative image of 5 animals per group. Bars are 50 µm.

### Succinate-induced anion secretion is carried by CFTR channels in human bronchial epithelial cells but succinate responses are impaired in human and mouse airway epithelium affected by the ΔF508 CFTR mutation, but recovered after ETI treatment of human ΔF508 cells

The contribution of CFTR to electrogenic anion secretion in mouse airways is debated, but cultured human airway epithelial cells show strong CFTR-dependent anion currents, and are the most suitable models to study CFTR function ^32^. In non-CF HBECs we found that succinate induced a negative change in I_SC_ that was prevented when cells were pre-incubated with the CFTR inhibitor CFTR_inh_172, but remained unaltered by TRPM5 or M3 inhibition (Figure 4A). These data indicate that succinate-induced anion secretion was CFTR-dependent but, importantly, that it was activated by a different signalling pathway than in mouse ^25^. We further confirmed the involvement of CFTR by performing the same experiment in cells isolated from CF patients homozygous for the ΔF508-CFTR mutation, and no changes in I_SC_ in response to succinate were observed in HBECs of this genotype (Figure 4A). It was previously shown that *SUCNR1* gene expression was decreased in HBECs from CF individuals ^25^, nevertheless we observed no changes in *SUCNR1* that might explain the absence of succinate-induced I_SC_ in CF HBECs (Figure S7A). Additionally, we found that gene expression of the Na^+^ coupled succinate co-transporter *SLC13A2* was not different between the two groups, indicating that (i) the absence of succinate-induced I_SC_ was not due to altered succinate-coupled Na^+^ absorption, or (ii) that, SLC13A2 activity contributed to succinate-induced I_SC_ in non-CF HBECs (Supplementary Figure 7A). Finally, non-CF and CF HBECs expressed equal amount of *TRPM5* and muscarinic receptors M1 and M3, and non-CF HBECs responded to the muscarinic agonist carbachol, confirming that TRPM5 and muscarinic signalling was not coupled to SUCNR1 activation in human airway epithelium (Figure S7A-B).

**Figure 4.**
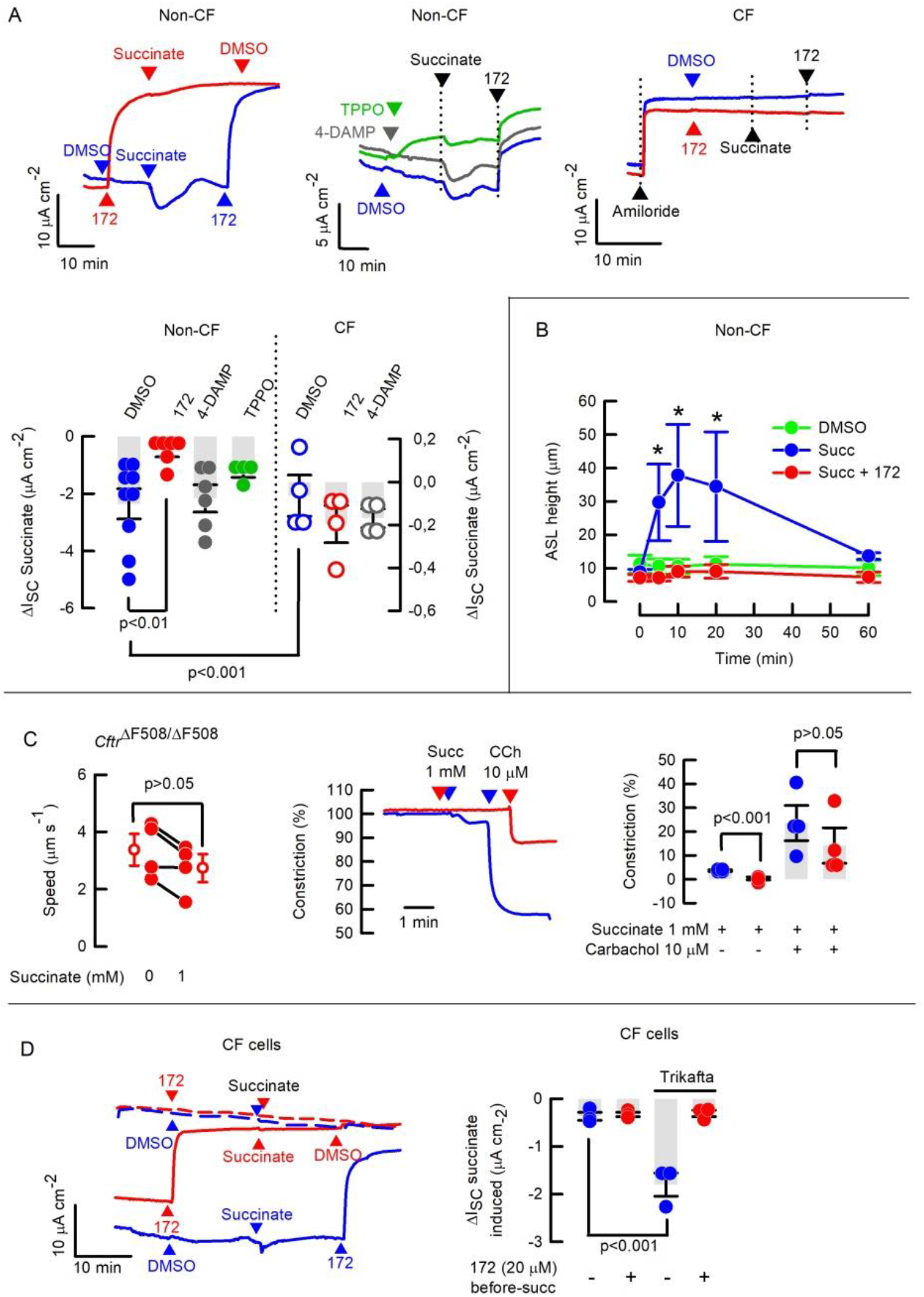
Succinate-induced anion secretion, mucus clearance and bronchoconstriction are impeded after CFTR inhibition or by mutations in CFTR, and recovered after ETI treatment in HBECS. (**A**) Representative traces of succinate induced I_SC_ in the presence or absence of 20 µM CFTR_inh_172 (172) or vehicle (DMSO), or in the presence of 30 nM 4-DAMP or 100 µM TPPO in non-CF HBECs. CF HBECs were treated with 172 before and after succinate. Summary of succinate induced ΔI_SC_ in non-CF and CF HBECs, bars are mean ± S.E.M and individual experiments are also included; n=4-9. ANOVA on Ranks. Succinate ΔI_SC_ in non-CF vs CF cells tested using Rank-sum test. (**B**) ASL height changes induced by 1 mM succinate were determined in the presence or absence of 20 µM 172, values are Mean ± S.E.M; n=6 for each condition, * indicates p<0.5; ANOVA on Ranks. (**C**) Effect of succinate effect on mucus transport measured in tracheas of *Cftr*^ΔF508/ΔF508^ mice. Rank sum test and representative traces of succinate induced bronchoconstriction in WT (blue line) and *Cftr*^ΔF508/ΔF508^ tracheal rings and Summary of bronchoconstriction experiments, Bars are Mean ± S.E.M, n=4 for each group, Rank-sum test. (**D**) Representative I_SC_ traces from cells treated with ETI (solid lines) or left untreated (dashed lines). Succinate-induced I_SC_ was measured in the presence of 20 µM Cftr_inh_172 (172) or vehicle (DMSO). And summary of ΔI_SC_ showing Mean ± S.E.M. and individual values determined in the presence or absence of 172. n=3 all groups, ANOVA on Ranks.

Consistent with a succinate-induced increase in CFTR-dependent anion secretion, succinate also increased ASL height, and this increase was also prevented by CFTR inhibition (Figure 4B). The effect of succinate on ASL height was not due to an osmotic effect since equimolar M-M-succinate has no effect on ASL-height (Supplemental Figure 7C).

To test if CFTR was necessary to initiate other succinate-induced functions, (e.g. mucus transport and bronchoconstriction) we used *Cftr*^ΔF508/ΔF508^ mice. When succinate was added to tracheas isolated from *Cftr*^ΔF508/ΔF508^ mice, there was no change in mucus transport rates or, surprisingly, no induced bronchoconstriction (Figure 4C). This last result, lead us to question whether M3 receptors were expressed in *Cftr*^ΔF508/ΔF508^ tissues, and as shown in Supplemental Figure 8 a strong signal for M3 receptors was observed in CF epithelial cells.

It was previously proposed that *SUCNR1* expression may be reduced in human-CF HBECs ^33^. So, we investigated whether ETI treatment recovered SUCNR1-dependent CFTR activation. As observed in Figure 4D ETI effectively restored succinate-induced anion secretion in human ΔF508-CF HBEC, this result is in accordance with the observation that in our CF HBECs, SUCNR1 gene expression was comparable to that of non-CF HBECs (Supplemental Figure 7A)

## DISCUSSION

MCC is an important epithelial innate defence mechanisms, that protects the lungs from infections. In this study, we presented substantial new evidence which demonstrated that activation of SUCNR1 initiated a signalling cascade to trigger airway *flushing* by coordinating anion secretion, mucus transport and mild bronchoconstriction in murine airways. We also reported that anion secretion was activated by succinate in HBECs and, most importantly, when CFTR was absent or mutated, succinate was unable to exert these beneficial functions. Taken together, these data point to the existence of a hitherto unrecognized signalling pathway for CFTR activation. The novel finding that CFTR can be activated through SUCNR1 is even more exciting as the expression of the receptor has been reported to occur in different cell types: brush cells in the mouse and PNEC in the human ^17,18,25^, suggesting an important role for SUCNR1 in CFTR activation that is conserved across species.

Recent evidence demonstrated that SUCNR1 activation increases cilia beating and anion secretion in mouse tracheas ^21^. However our results provide important new information about the underlying molecular mechanisms of this process. First, our data demonstrate that SUCNR1 activates HCO_3_^-^ secretion. This is of particular importance as we, and others, have clearly demonstrated that HCO_3_^-^ secretion is essential for airway homeostasis, particularly for mucus function and bacterial killing ^30,34,35^. Second, mucus transport is enhanced by the SUCNR1-dependent increase in anion secretion. Third, bronchoconstriction couples to SUCNR1 activation. Bronchoconstriction is expected to reduce the lumen surface, to increase ASL height further than just by fluid secretion alone, driving mucus and captured particles out of the airways more efficiently. Even though bronchoconstriction can be associated with pathological airway-flow restriction ^36^, we calculated that the reduction of airway flow after succinate to be less than 20%, which is considered to be physiologically mild bronchoconstriction in humans ^37^.

The existence of SUCNR1-induced HCO_3_^-^ secretion in the mouse airways is further strengthened by the finding that M3 receptors are co-expressed with the SLC4A4 co-transporter in CCSP+ secretory cells from the mouse trachea, that also express CFTR and TMEM16A channels ^38^. This direct coupling of cholinergic signalling with anion secretion may act as the first event that activates cilia beating through the increase of ASL volume ^29^, and further potentiated by alkalization of the ASL that has been shown to increase cilia beating in other models ^39^. The proposal that succinate activates airway cleaning by the spread of Ca^2+^-signalling through connexins ^21^ is questioned by our data, as succinate was unable to increase mucus clearance in *Cftr*^ΔF508/ΔF508^ mice, suggesting that CFTR activity is upstream in the succinate-induced airway clearance activation pathway.

Increased succinate levels have been found in the airways of CF individuals and CF-HBECs ^27^. These observations prompted us to study whether SUCNR1 signalling was CFTR-dependent, and moreover, if it was also altered in CF models. First, we found that succinate-induced anion secretion and subsequent increase in ASL height were fully blocked by pre-treatment with CFTR_inh_172 in non-CF HBECs. Second, succinate-induced anion secretion was absent in HBECs obtained from CF individuals. In addition, we observed that succinate-induced anion secretion was TRPM5 and M3-independent, which is in agreement with previous observations that point towards an SUCNR1-ATP-P2Y1 axis in human and pig submucosal glands instead of SUCNR1-acetylcholine-M3 axis seen in murine trachea ^21,25^.

Mice have been overtly discredited as a *bona fide* CF model, due to the absence of an obvious CF-phenotype in the lungs and difficulties to measure CFTR currents ^40^. While in humans Cl^-^ is transported out of the cells preferentially by CFTR, the identity of the channels underlying Cl^-^ secretion across the mouse airways is still an open question. Nevertheless, *Pseudomonas aeruginosa* clearance studies have clearly demonstrated that the CF-mouse is defective in clearing this bacteria, indicating a CFTR-dependence for such function ^41,42^. In agreement with this evidence, we observed that mucus transport was not increased after succinate addition in *Cftr*^ΔF508/ΔF508^ mice, confirming that the model is affected by deficient MCC.

Even though we did not directly measure HCO_3_^-^ secretion HBECs in this study, we and others, have previously shown that CFTR activation in human airway cells is an essential step for HCO_3_^-^ secretion which increases ASL pH by activation of SLC4A4 and SLC24A4-Pendrin ^30,43,44^. HBECs do not contain airway smooth muscle and hence, cannot be used for bronchoconstriction studies, and we were unable to test SUCNR1-dependent function. Nevertheless, analogous to our bronchoconstriction data, pig submucosal glands contract when myoepithelial cells are stimulated by the SUCNR1-ATP-P2Y1 axis, indicating that succinate leads to activation of contractible (myoepithelial or smooth muscle) cells surrounding different airway epithelial structures ^25^.

Succinate can be released from inflammatory macrophages to activate airway *flushing* as a physiological response, but the accumulation of succinate in the airways promotes *Pseudomonas aeruginosa* colonization in the airways of CF-patients ^27^. In this scenario, loss of SUCNR1-induced *flushing* could therefore contribute to exacerbations in CF patients. Our data shows that restoration of CFTR function following ETI treatment was accompanied by recovery of SUCNR1-induced anion secretion, which would be expected to aid the desired correction of innate immune functions of the airways. For example, SUCNR1 is also found in CFTR expressing cells, such as macrophages, whose antimicrobial activity against *Pseudomonas aeruginosa* is recovered after ETI in CF-individuals ^45^. Even though, the function of SUCNR1 has not being studied in CF macrophages, these cells display an inflammatory phenotype after SUCNR1 knock-down, suggesting that the correction of the SUCNR1-CFTR axis in macrophages might also help to ameliorate inflammation ^46^. Finally, loss of SUCNR1 induces hypoinsulinaemia and hyperglycaemia in mice, suggesting that the loss of SUCNR1-CFTR signalling might favour the development of diabetes in CF-patients ^47^.

Our study has some limitations?. Despite differences in expression and downstream signalling we clearly show that succinate-SUCNR1 signalling leads to CFTR activation in both species, dependent on SUCNR1 activation. Nevertheless, experimental evidence from human or *ex-vivo* human tissues showing succinate responses and the clearance of pathogens from the *Sucnr1*^-/-^ mouse or from a specific *Sucnr1* knock down from airway epithelium are still lacking.

In summary our data describes a new signalling pathway that activates CFTR through the SUCNR1 in both human and mouse airway epithelial cells. We propose that chemosensing of succinate via SUCNR1 activation form an important part of the innate defence mechanism of the airways, by flushing out mucus and pathogens. The impairment of the succinate-SUCNR1-CFTR axis in human diseases where CFTR activity is greatly decreased like CF and COPD, might therefore contribute to disease worsening, but at least in the case of CF, this should be corrected upon restoration of CFTR function.

## MATERIAL AND METHODS

### Reagents

The CFTR modulators tezacaftor (CAS1152311-62-0), ivacaftor (CAS873054-44-5) and elexacaftor (CAS2216712-66-0) were purchased from Insight Biotechnology Limited. Other chemicals, including succinic acid (224731), mono-Methyl hydrogen succinate (M81101), triphenylphosphine oxide (T84603), carbamoylcholine chloride (C4382), amiloride hydrochloride hydrate (A7410), adenosine 5’-triphosphate disodium salt hydrate (A7699), uridine 5’-triphosphate disodium salt hydrate (U6750), fumarate (F1506), itaconate (I29204), malate (M1296), denatonium (D5765), ANI9 (SML1813) and S0859 (SML0638) were of reagent grade and supplied by Sigma-Aldrich UK Ltd. CFTRinh-172 (3430), Pyrenzipine (1071) and 4-DAMP (0482) were from Tocris Biosciences. U73122 was from Merck (112648-68-7) and eserine from MedChemExpress (HY-N6608). To achieve final concentrations, stock solutions were diluted in Krebs Ringer Buffer (KRB), and DMSO, except S0859 which was re-suspended in ethanol. DMSO and ethanol concentrations were maintained at below 0.1% so as not to alter epithelial properties.

### Animals

The wild type C57BL/6 J mice were obtained from The Jackson Laboratories (USA). the *Cftr*^tm1Eur^ mouse (named here as *Cftr*^ΔF508/ΔF508^) was kindly donated by Dr B. Scholte ^48^. The *Sucnr1*^-/-^ mouse line was generated at the Centro de Estudios Científicos (CECs), as described below. Animals were bred and maintained in the Specific Pathogen Free mouse facility of CECs with access to food and water ad libitum. Six–10 week-old male or female mice were used. Sex was not considered as a biological variable. The *Slc4a4*^-/-^ mouse line was kindly donated by Dr. Gary Shull, bred in the original 129S6/SvEv/Black Swiss background ^49^ and used for experiments at 14 days old. Unless otherwise stated, all procedures were performed after mice were deeply anesthetized via i.p. injection of 120 mg/kg ketamine and 16 mg/kg xylazine followed by exsanguination. All experimental procedures were approved by the Centro de Estudios Científicos (CECs) Institutional Animal Care and Use Committee (#2022–04) and are in accordance with relevant guidelines and regulations. Lung tissue samples of the *Trmp5*^egfp/+^ mice were obtained from mice generated by Clapp and co-workers ^50^.

### Generation of the *Sucnr1*^-/-^ mouse line

This mouse was generated using CRISPR/Cas9 editing, where a couple of guides and Cas9 enzyme were used to delete part of exon 2 of *Sucnr1* gene. Specific guides located at the beginning of exon 2 and close to the stop codon were designed using the CRISPOR tool (http://crispor.tefor.net/) ^51^. The best score guides were selected and synthesized as crRNA at IDT DNA (USA) together with the universal scaffold, tracrRNA. The S.p. HiFi Cas9 protein, crRNAs and tracrRNA were electroporated in zygotes obtained from females C57BL/6 and then transferred to a pseudopregnant female (B6CBA). Genotyping was performed by PCR from genomic DNA extracted from pup tail biopsies with primers For1 and Rev1. The PCR products were sequenced to determine the exact size of the genomic DNA deletion (See Suppl. Figure 1).

### Ussing chamber experiments

Mouse tracheae were placed in P2306B of 0.057 cm^2^ tissue holders and placed in Ussing chambers (Physiologic Instruments Inc, San Diego, CA, USA). Tissues were bathed with bicarbonate-buffered Ussing solution (pH 7.4) of the following composition (in mM): 120 NaCl, 25 NaHCO_3_, 3.3 KH_2_PO_4_, 0.8 K_2_HPO_4_, 1.2 MgCl_2_, 1.2 CaCl_2_ or HEPES buffer: 130 NaCl, 5 KCl, 1 MgCl_2_, 1 CaCl_2_, 10 Na-HEPES (pH adjusted to 7.4 using HCl); supplemented with 10 D-Glucose, gassed with 5% CO_2_–95% O_2_ (bicarbonate buffer) or 100% O_2_ (HEPES buffer) and kept at 37 °C. The transepithelial potential difference referred to the serosal side was measured using a VCC MC2 amplifier (Physiologic Instruments Inc). The short-circuit currents were calculated using the Ohm’s law as previously described ^30^. Briefly Ca^2+^-dependent anion secretion was induced by 1 mM succinate, 50 µM carbachol and 50 µM UTP. The ΔIsc values were calculated as the area under the curve (A.U.C.) of the first 2 min post stimulation using the Acquire & Analyze 2.3 v software. Tissues with transepithelial resistance (R_te_) values below 50 Ωcm^2^ were discarded as they were not suitable for *bona fide* electrophysiological determinations ^52^.

### Particle track and mucus speed measurements

We used the previously published methodology which originally used piglet tracheas, and adapted this for mouse tracheas ^53^. Mice were euthanized with Isoflurane (Baxter healthcare corporation), and dissections were performed on the heart, lungs, and trachea, with a particular focus on precise dissection of the thyroid cartilage. Tissues were maintained in cold Ussing solution gassed with 5% CO_2_ until use. Each trachea was meticulously cleaned and mounted on a 35 mm diameter plate coated with a layer of Sylgard 184 (The Dow Chemical Company). Then the lungs were pinned, and small incisions were made to release air, preventing the formation of bubbles that could interfere with measurements. Tracheas were stained with a 0.0125% alcian blue solution diluted in Ussing solution for 5 minutes at 37°C and then mounted on an horizontal chamber for PTS measurements ^30^, or in an inclined platform at a 30° angle for mucus transport, with temperature maintained between 28°C and 32°C using a temperature controller. Finally images were acquired at 2-second intervals for approximately 10 minutes using a Motic camera (Model Plus 1080X) and the mucus movement was analyzed in the image sequence using NIH Fiji Image J software, utilizing the TrackMate plugin. Videos were generated using Movavi software (Supplemental Fig 4A).

### Bronchoconstriction studies

We used the precision cut lung slice (PCLS) technique ^54^, adapted for mouse trachea. Animals were euthanized with a lethal dose of anesthesia (ketamine (100mg/ml) / xylazine (10mg/kg)), and then the trachea and bronchial tree were insufflated with 2% low-melt agarose (Agarosa Lafken LM). Subsequently, the trachea and main bronchi were removed, cut in a McIlwain™ Tissue Chopper into 150-200 μm slices. Tissues were incubated at 37°C in a HCO_3_^-^ buffered solution for 15 minutes, to then be washed two to three times to eliminate remaining agarose. Subsequently, the slices were mounted in a continuous flow chamber under a video-microscope, and perfused (1ml min^-1^) with 50mM KCL for pre-contraction, returned to the HCO_3_^-^ buffered solution and then exposed to succinate (1mM) and carbachol (10µM). Photos were taken every 2s for a total of 15 to 20 min. Subsequently, the internal area of the trachea was measured using ImageJ NIH and changes in area over time calculated after normalizing to the initial area of the trachea. Videos were generated using Movavi software. Differences in the airway flow for each experiment, was determined using the Poiseuille equation: Q=π·ΔP·r^4^/8·η·L. The value of the change in the radius (r) was obtained using the expression for the calculation of the area of a circle: A=π·r^2^, before and after succinate addition.

### Immunofluorescence

Mice were euthanized by exsanguination by severing the inferior vena cava under deep anesthesia. Mice were perfused with 10% formalin/PBS. Trachea and lung were removed and incubated overnight in 10% formalin/PBS at 4 °C. Paraffin sections (5 μm) were treated with 1X EDTA buffer pH 8.0 (Diagnostic Biosystem), blocked with 2.5% normal goat serum (Vector Laboratories cat# S-1012), and incubated with 1:100 anti-CHRM3 (anti-M3r; Alomone, AMR-006,) 4 °C overnight. Sections were incubated with secondary antibody 1:1,000 Alexa fluor 488 goat anti Rabbit IgG (H+L) (Invitrogen, cat# A-11008) 1 hr at room temperature. For colocalization 1:100 anti-CHRM3 was incubated with 1:500 anti-Clara Cell Secretory Protein (CCSP; Merck Millipore cat#07– 623), 1:200 anti-alpha tubulin (Santa Cruz, cat#sc-5286) or 1:100 anti-NBCe1 (Alomone, ANT-075,) overnight at 4 °C, and incubated with secondary antibody 1:1,000 Alexa fluor 555 goat anti Rabbit IgG (H+L) (Life Technologies, cat#A-A21428), 1:1,000 Alexa fluor 568 goat anti Mouse IgG2b (H+L) (Life Technologies, cat#A-A21144) or 1:1,000 Alexa fluor 555 goat anti Rabbit IgG (H+L) (Life Technologies, cat#A-A21428) respectively. Detection of SUCNR1 was performed using the anti-GPR91 (SUCNR1; Alomone #ASR-090) and TRPM5 with anti-TRPM5 (Alomone, #ACC-054), both tested with and without antigen retrieval using pH 6.0 or pH 8.0 buffers and overnight incubation at RT or 4 C°. Nuclei were stained with 1:2,000 propidium iodide (Invitrogen, cat#P21493). All immunofluorescence images were captured using a confocal microscope (Olympus FV1000).

### Human bronchial cells culture and electrophysiology

#### Cell culture

Primary non-CF (n = 1 donor) and CF (n = 2 donors, all F580del/F508del genotype) Human Bronchial Airway Epithelial Cells (HBECs) were a gift from S. H. Randell (University of North Carolina (UNC) CF Center Tissue Procurement and Cell Culture Core). They were obtained under protocol #03-1396 approved by the UNC Biomedical Institutional Review Board. HBECs were expanded using the conditionally reprogrammed cell (CRC) culture method ^55^. In brief, HBECs were seeded onto modified 3T3J2 fibroblasts (inactivated with mitomycin C 4 µg/ml, 2 h, 37 °C) and grown in medium containing the ROCK inhibitor Y-27632 (10 μM) until 80% confluent. Cells then underwent double trypsinisation to first remove the fibroblasts and then detach the HBECs. At this stage, the HBECS were counted and cryopreserved for future use. After thawing, HBECs were seeded directly onto human placental collagen (HPC, Sigma-Aldrich, C8374)-coated permeable supports (Costar Transwells, 6.5 mm; 3470) in bilateral BEGM growth medium. After two days, when cells reached confluency, the medium was replaced with the commercially available differentiation medium StemCell PneumaCult™-ALI Medium (Catalog #05001). After three days with bilateral differentiation, the apical medium was removed and the HBECs fully differentiated under Air-Liquid Interface (ALI) conditions as described previously ^56^. Ciliogenesis started approximately 12-15 days after seeding and the cells were used for experiments between days 25 and 35 after seeding. Some cells were also pre-treated with a combination of elexacaftor (VX-445, 2 µM), tezacaftor (VX-661, 3 µM) and ivacaftor (VX-770, 1 µM) (ETI) for 24 h at 37 °C, before experiments.

#### Gene expression analysis

The relative gene expression between non-CF and CF samples was obtained from a RNAseq dataset of the same cells accessible through Gene Expression Omnibus (GEO series accession number GSE154905) as described in Saint-Criq et al ^56^.

#### Short-circuit current measurements in Ussing chambers

Cells grown on Costar 6.5 mm inserts were mounted in the EasyMount Ussing Chamber System (VCC MC8, Physiologic Instrument Inc, USA) and bathed bilaterally in a HCO^3-^ KRB and continuously gassed with 95% O2-5% CO2 and maintained at 37 °C. The KRB solution contained (in mM) 115 NaCl, 25 NaHCO_3_, 5 KCl, 1.0 MgCl_2_, 1.0 CaCl_2_, 5 Glucose (pH 7.4). Epithelia were voltage-clamped at 0 mV and left to equilibrate for at least 20 min before ion transport studies were performed. The general protocol consisted of sequential and cumulative addition of compounds as described in the result section. All compounds were added to the solution bathing the apical membrane only. The transepithelial short-circuit current (I_sc_) was recorded using Ag-AgCl electrodes in 3 M KCl agar bridges and results expressed as µA.cm^−2^ and analysed using Acquire & Analyze software (Physiologic Instruments) as previously described ^57^.

#### Human bronchial cells culture and ASL height

Primary non-CF cultures (3 donors) were mucosally washed 3 x in PBS with the last wash extended for 30 min at 37°C in a 5% CO2 incubator. After aspiration of all PBS, 0.5 mg/ml rhodamine-dextran (Life Technologies, 10 kDa, neutral) in 20 µl Ringer’s solution was added mucosally. Excess dye was aspirated from the mucosal surface by Pasteur pipette leaving residual dye that was ∼7 µm in height. Cultures were incubated for an additional 30 min and then ASL height was measured. Cells were placed in Ringer’s solution during imaging with perfluorocarbon (FC-770, 3M FluorinertTM) added mucosally to prevent ASL evaporation. 20 images per culture were obtained automatically at pre-determined pointsby XZ confocal microscopy using a Leica SP8 microscope with a 63X glycerol immersion objective. To prevent bias, images were analyzed automatically using a macro written in ImageJ software (National Institutes of Health freeware) to determine ASL heights.

## Acknowledgements

Funded by FONDECYT 1221257, and FOVI 220118 (C.A.F.); Cystic Fibrosis Trust, SRC013 (M.A.G); FONDECYT 1211838 (M.A.C.); Grant in aid from Boehringer Ingelheim (C.W); NIH NIDCD DC000566 (D.R.); Vicerrectoría de Investigación y Doctorados de la Universidad San Sebastián, Chile – USS-FIN-23-FAPE-14 (I.C.); FONDECYT 3220672 (S.V.); TARRAN22G0 (RT). We thank Dr. Joe Wrennall for writing the Image J ASL height macro.

---

**Figure S1.**
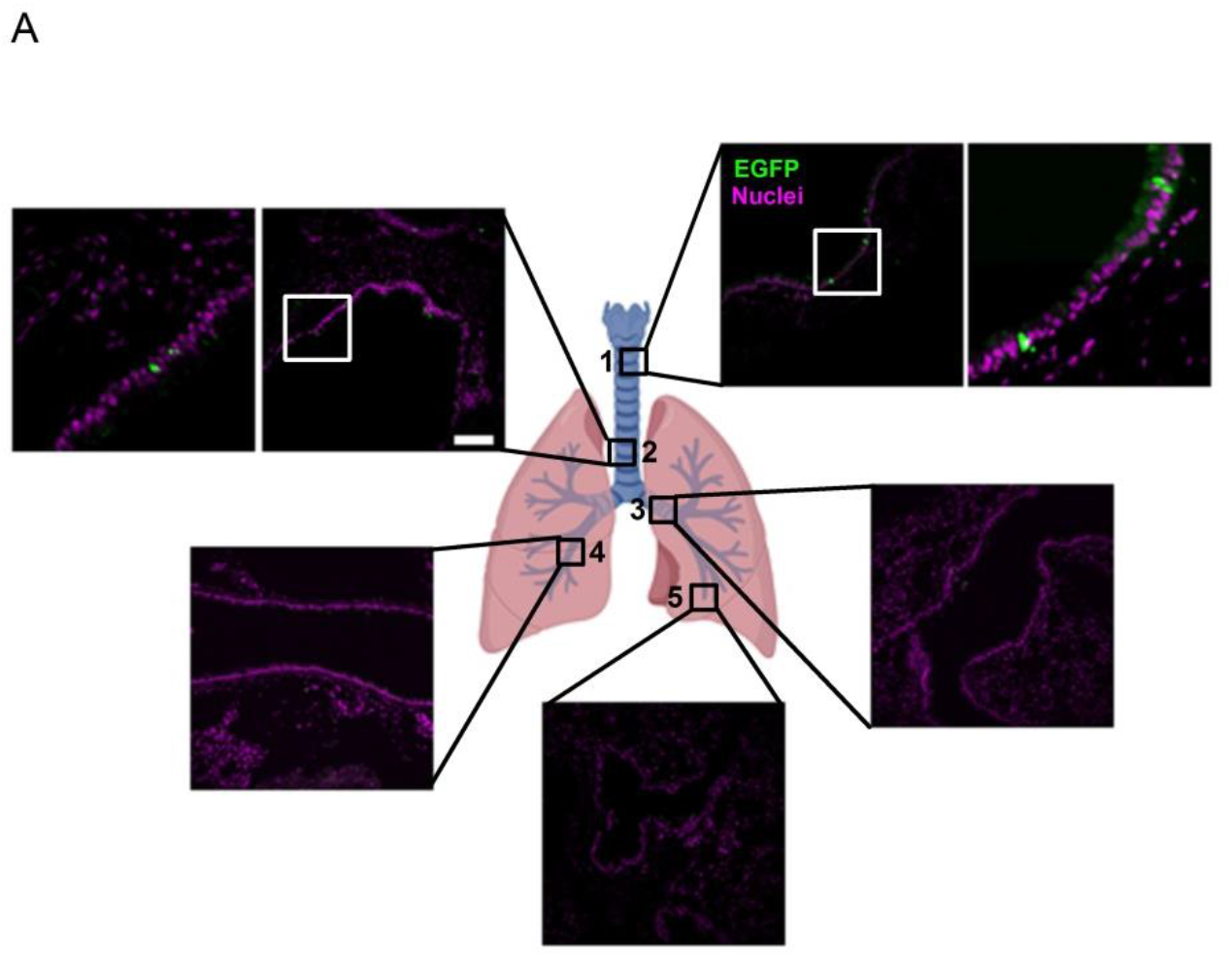
TRPM5 positive cells are present in the proximal airways of the mouse. **(A)** Immunodetection of EGFP in TRPM5-EGFP mouse tissues. Numbers indicate 1. Upper trachea; 2 Lower trachea; 3 main bronchi; 4 bronchi and 5 terminal bronchiole. Representative images of 3 different animals. Bar = 50µm.

**Figure S2.**
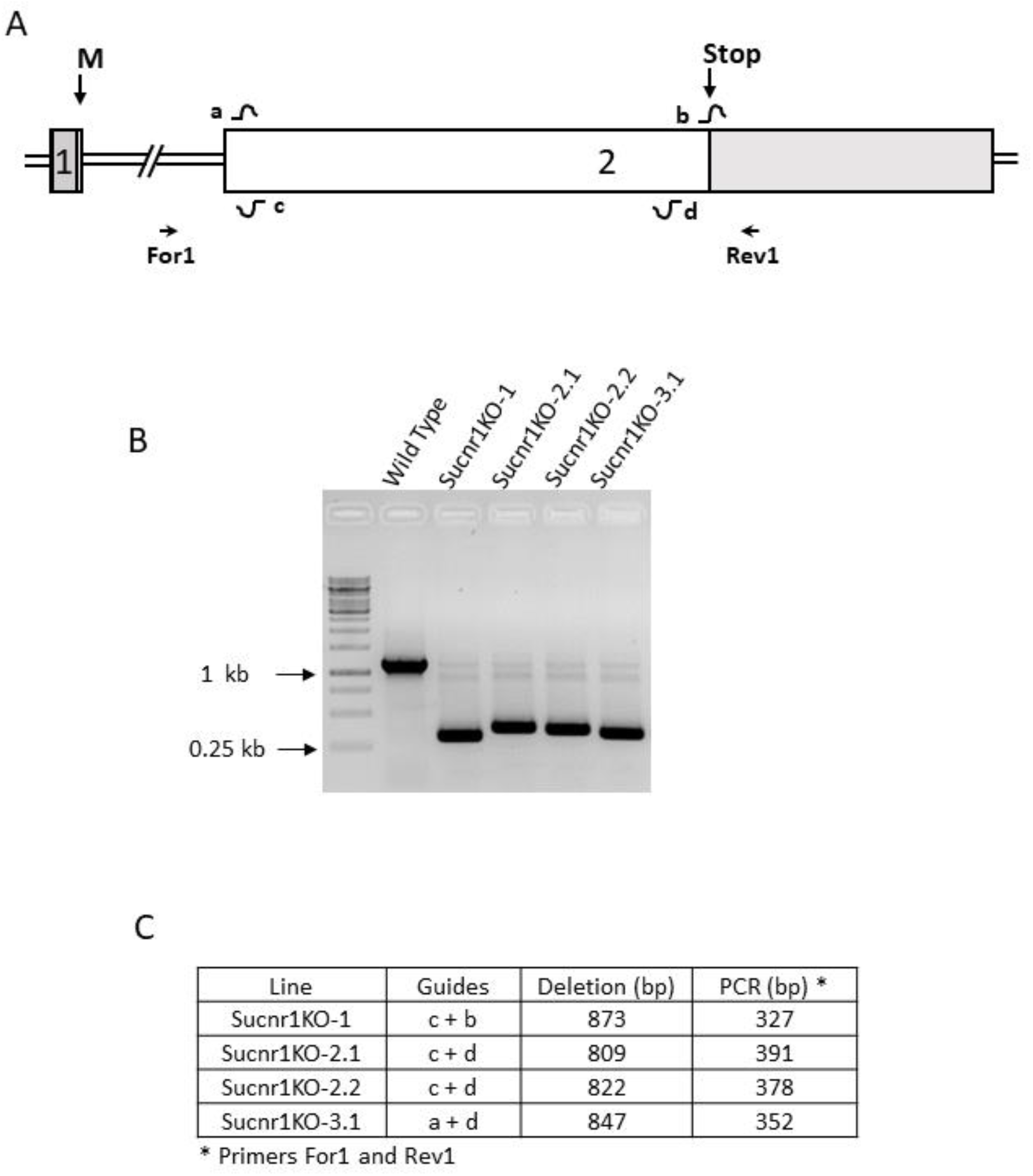
Strategy for the generation of the *Sucnr1*^-/-^ mice. **(A)** Scheme of the Sucnr1 gene showing its two exons (boxes 1 and 2), the coding regions in white and methionine (M) and stop codon (Stop) with down arrows. The guides are represented as a, b, c and d and the primers For1 and Rev1 for genotypification with right and left arrows. **(B)** Agarose gel showing the PCR products of WT and Sucnr1KO lines obtained using the primers For1 and Rev1. The samples used for each line showed belong to heterozygous animals. **(C)** Table to show the couple of guides used to generate the Sucnr1KO lines, the size of the deletion of exon 2 obtained after edition and the PCR product size observed with primers Fo1 and Rev1. The sequences of guides and primers are: tgagaattggttggcaacag (+1) b: tcaagtcccttacatccttc (+1) c: cgattgcataaaatgcagag (−1) d: ctctgtaatggtctcccatg (−1), For1: tcgtattgaaggtccatgcc and Rev1: gctaaagcctagaaatggtgg.

**Figure S3.**
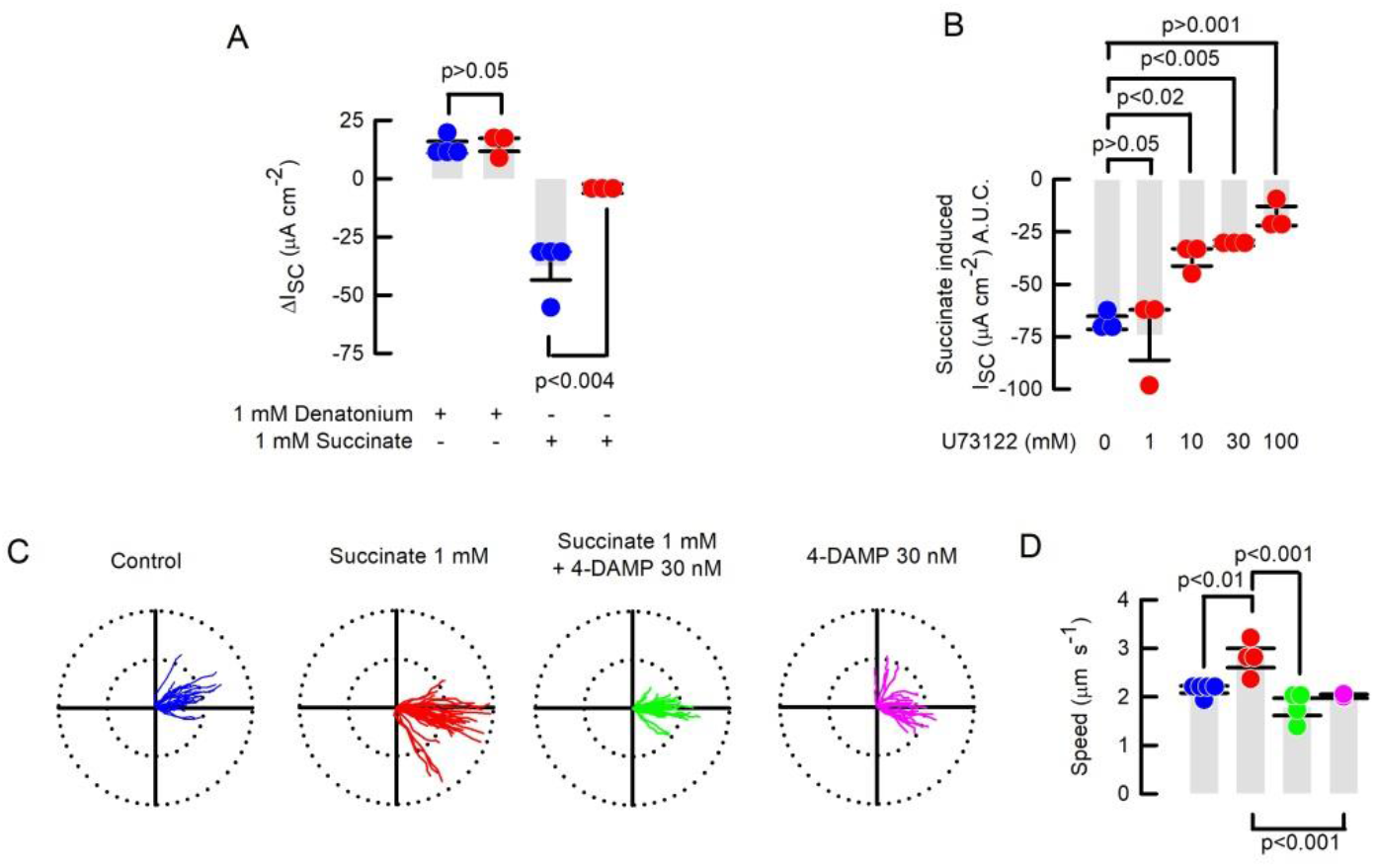
Anion secretion and increased particle track speed are impaired after *Sucnr1* silencing or signalling inhibition in mouse trachea. (**A**) Summary of ΔI_SC_ measurements in wild type (blue dots) or *Sucnr1*^-/-^ (red dots) mouse tracheas after addition of denatonium and succinate. Bars are mean ± S.E.M; n=4 and n=3 for wild type and *Sucnr1*^-/-^ respectively; Rank sum test. (**B**) Summary of succinate-induced I_SC_ measurements in wild type tracheas in the absence (blue dots) or the presence (red dots) of increasing concentrations of U73122. Bars correspond to mean ± S.E.M; n=3 for each group; ANOVA on ranks. (**C**) Beads tracking of PTS experiments performed in wild type tracheas in the presence or absence of succinate and/or 4-DAMP. Radius of the polar plots is 50 µm. (**D**) Summary of PTS experiments for speed. Bars indicate mean ± S.E.M.; n=4 for each group but n=3 for 4-DAMP alone; ANOVA on ranks.

**Figure S4.**
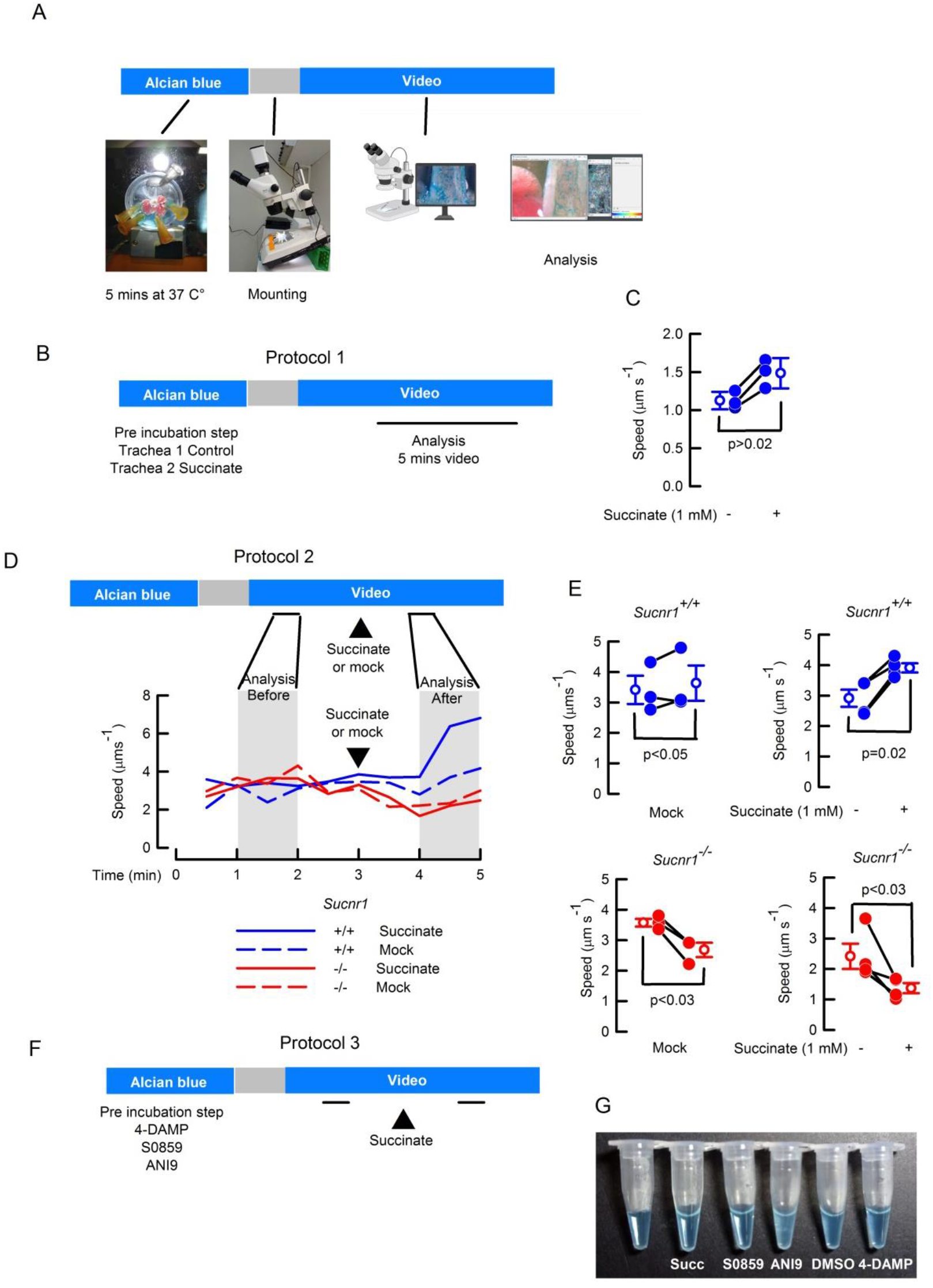
Set-up ad protocols for mucus transport studies in the mouse trachea. (**A**) Description of mucus transport set-up and protocol. More detailed information available in the corresponding Methods section. (**B**) Mucus transport protocol 1 using succinate pre incubation step parallel to mucus staining. (**C**) Succinate effect on mucus speed determined using protocol 1. Values are means ± S.E.M., n=4; Rank sum test. (**D**) Protocol 2 for mucus transport speed indicating times for data analysis after and before the addition of buffer (mock) or 1 mM succinate. The lower panel includes representative traces of experiments including wild type (*Sucnr1*^+/+^) or *Sucnr*^-/-^ tracheas after mock or succinate addition. (**E**) Summary of experiments obtained using protocol 2. Values are means ± S.E.M., n=3 for mock and n=4 for sucinnate; Rank sum test. (**F**) Protocol 3 for mucus transport including pre incubation step with different inhibitors. (**G**) Image showing the Alcian blue solution in buffer after the addition of different inhibitors and molecules. Note the cloudly aspect of the tube where ANI9 was added.

**Figure S5.**
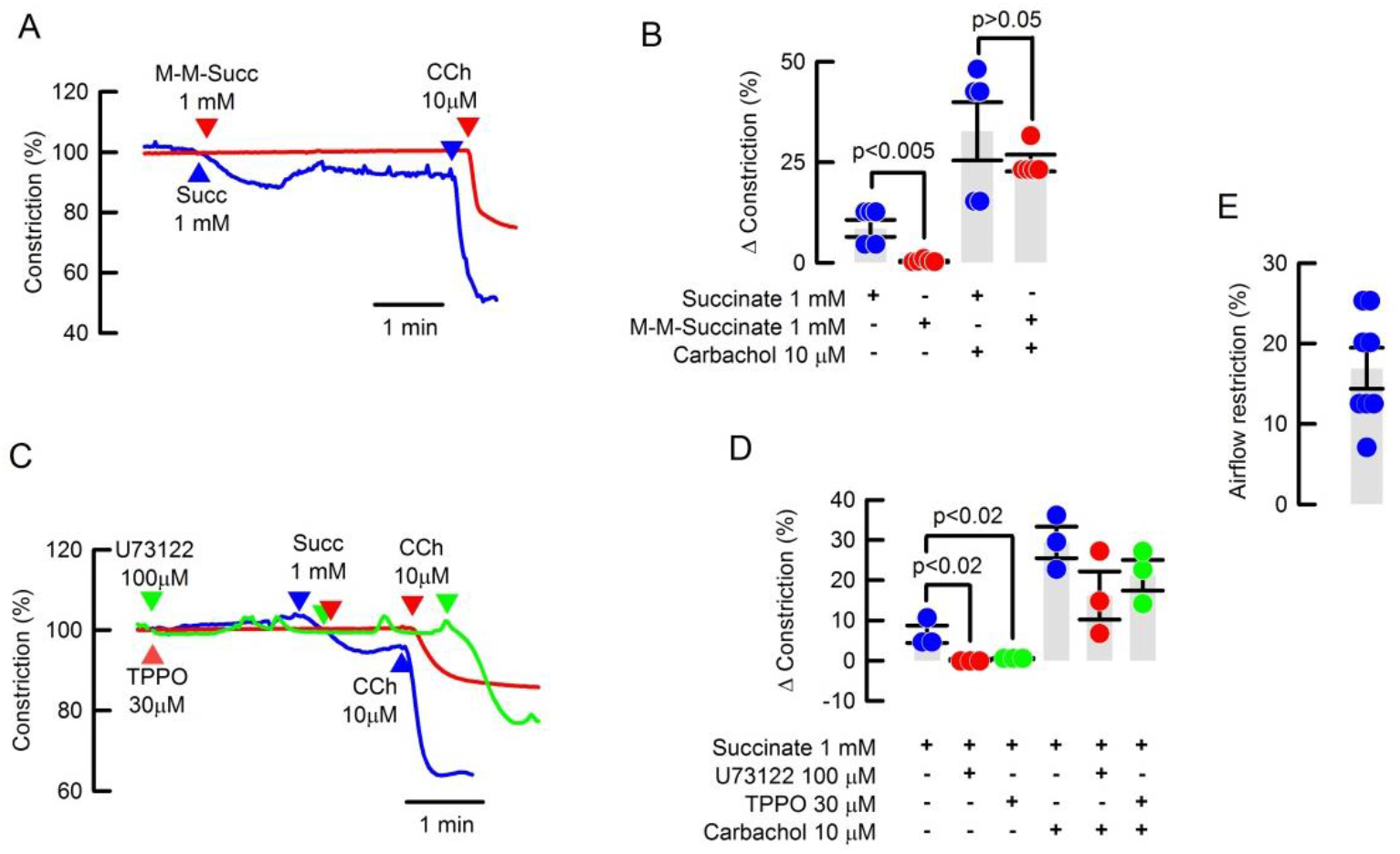
Succinate-induced bronchoconstriction is dependent on SUCNR1 and downstream signalling. (**A**) Representative traces for bronchoconstriction after 1 mM succinate or 1 mM M-M-succinate addition. (**B**) Summary of experiments as in (A) including CCh stimulation; n=5 for each gruop; Rank sum test. (**C**) Representative traces for succinate-induced bronchoconstriction in the presence of TPPO or U73122. (**D**) Summary of experiments as in (C) including CCh stimulation; values are means ± S.E.M.; n=3 for each group; ANOVA on Ranks. **(E)** Summary of calculated airflow restriction after succinate in the mouse trachea, value is mean ± S.E.M.; n=8.

**Figure S6.**
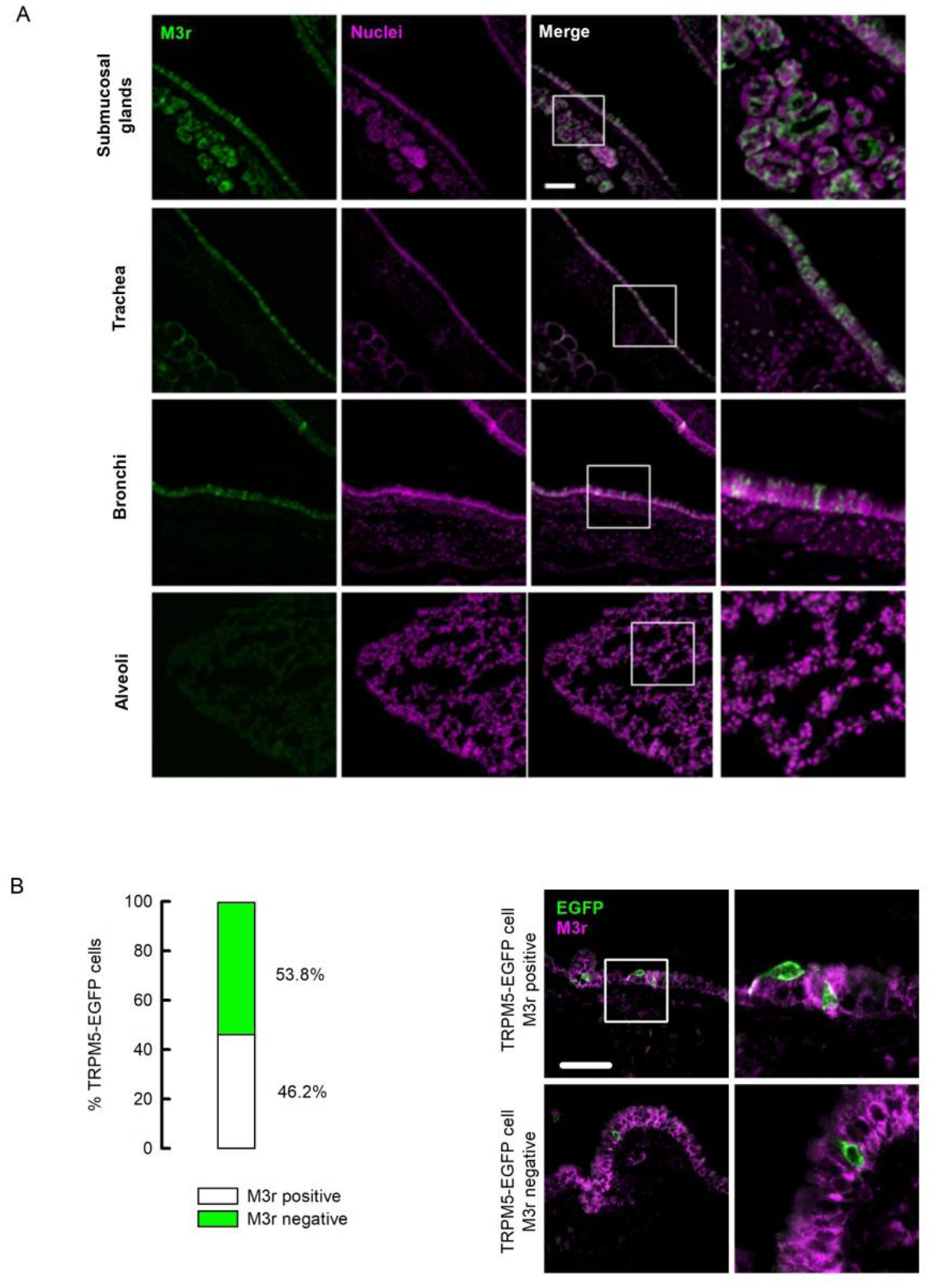
M3 receptor expression in airway epithelium and TRPM5+ cells in the mouse. (A) M3 receptors expression in the submucosal glands and proximal to distal airways in the mouse. Figures are representative of 4 different wild type animals. (B) Summary of the expression of M3 receptor in TRPM5-EGFP+ cells and representative images of the co-staining. Data is taken from 13 cells from 3 different animals.

**Figure S7.**
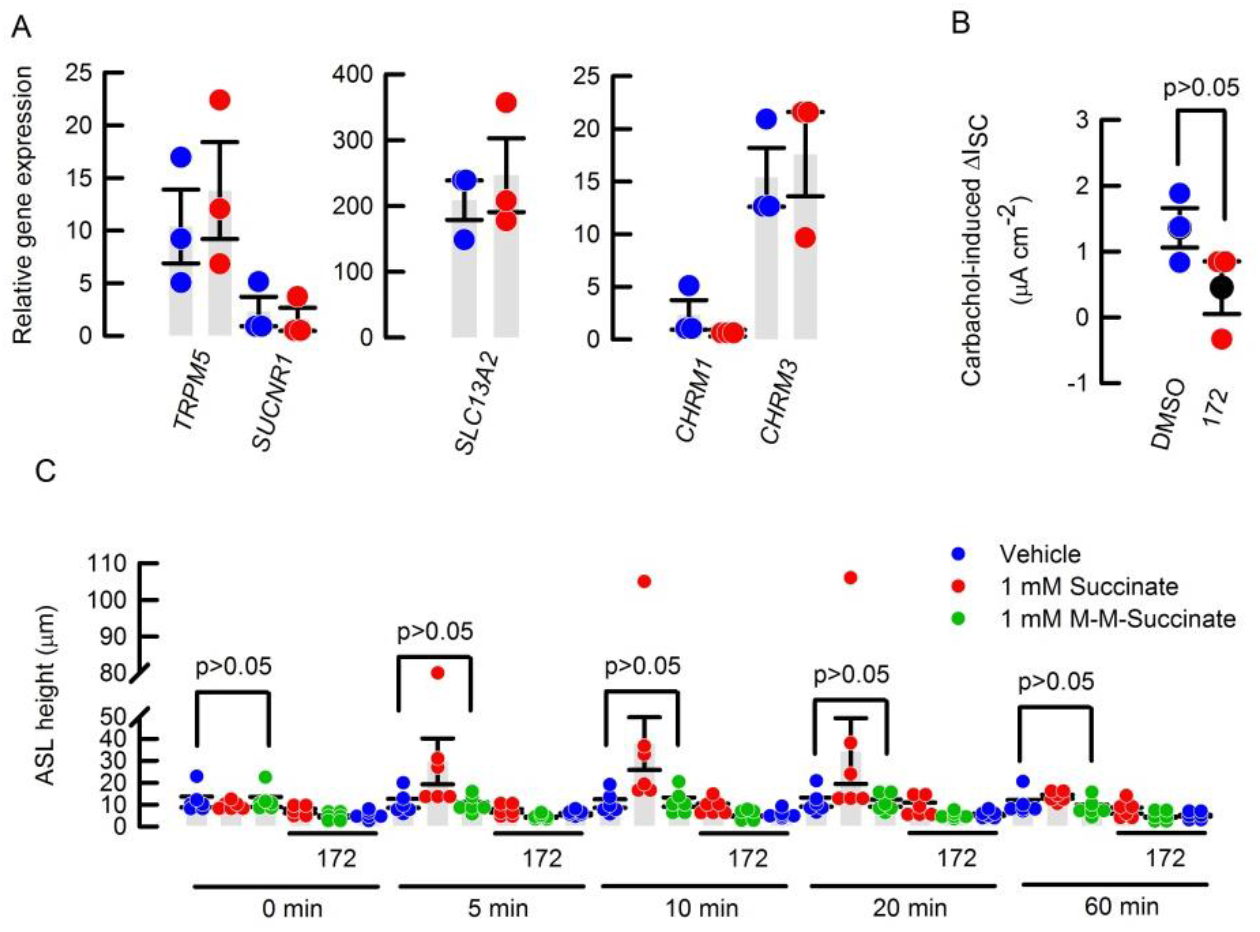
Gene expression, carbachol-induced I_SC_ and ASL height in HBEC. (A) Non CF (blue dots) and CF (red dots) HBEC gene expression. (B) Carbachol effect in HBEC pre-treated with vehicle (DMSO) or CFTR_inh_172 (172). Values are mean (black circle ± S.E.M.) Rank sum test. (C) ASL height changes induced by 1 mM succinate or 1 mM M-M-succinate were determined in the presence or absence of 20 µM 172, values are Mean ± S.E.M; n=6 for each condition.

**Figure S8.**
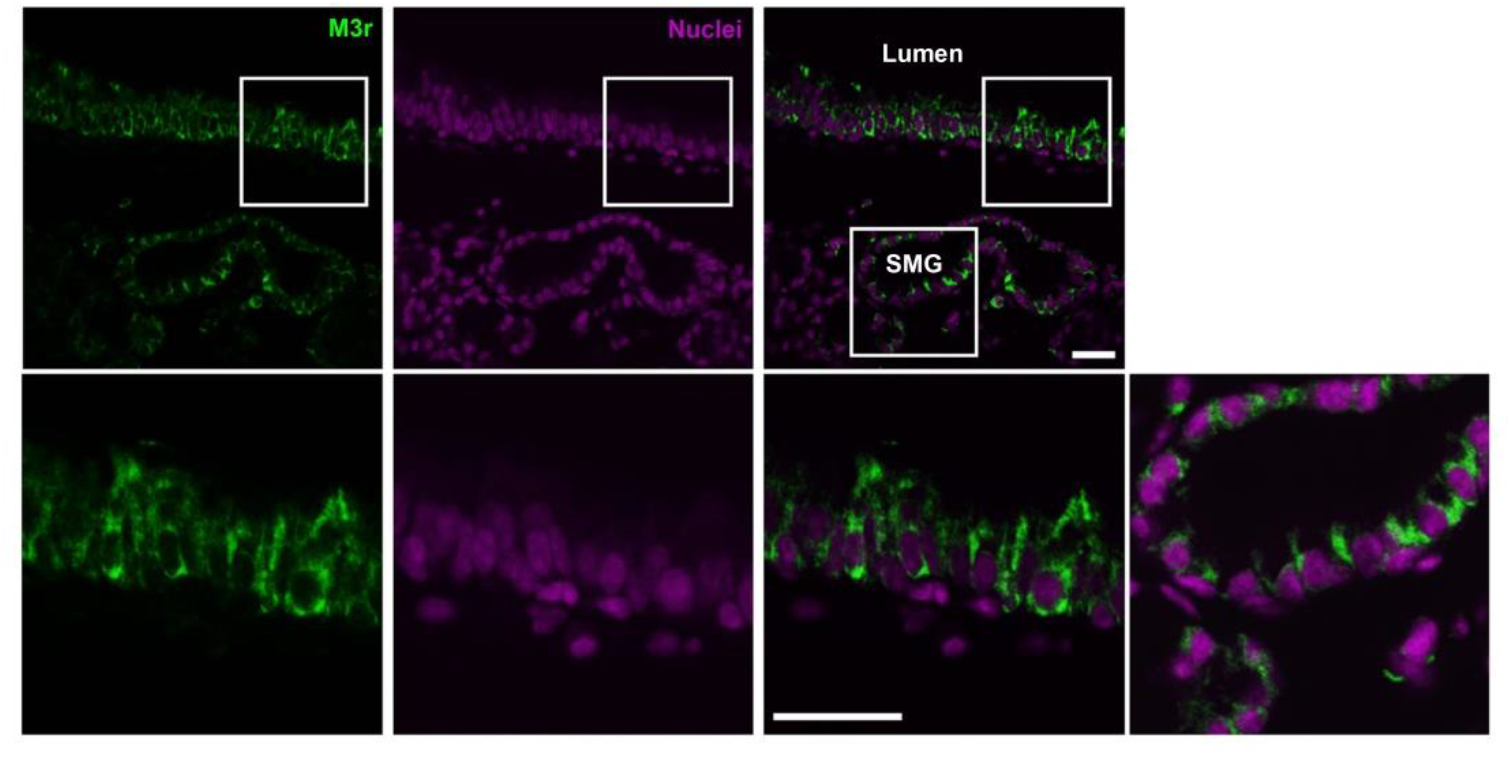
M3 receptor expression in the *Cftr*^ΔF508/ΔF508^ mouse airway epithelium. M3 receptors expression in the trachea and submucosal glands (SMG) of the *Cftr*^ΔF508/ΔF508^ mouse. Figures are representative of 3 different animals.

